# FastTomo: A SerialEM Script for Collecting Electron Tomography Data

**DOI:** 10.1101/2021.03.16.435675

**Authors:** Albert Xu, Chen Xu

## Abstract

FastTomo is a SerialEM script for collecting tilted specimen images in transmission electron microscopes to be further used in tomographic reconstruction. It achieves a speedup over conventional tracking methods by minimizing the usage of off-target tracking shots, and instead applies proportional control to the specimen images. Movement in the Z coordinate is estimated prior to each tilt series in a separate calibration routine. Overall, this method is fast and reliable when the field of view is at least 1 um, and can tolerate minor errors in setting eucentric height. The implemented tilt series schemes include the unidirectional, bidirectional, and dose-symmetric schemes.

## I. Introduction

Electron tomography is a method of reconstructing a 3D volume from 2D projected images taken at various angles by an electron microscope. In order to successfully collect a series of images, two major problems must be addressed. The first problem is that the sample will move out of view of the camera, so the specimen must be tracked in X and Y coordinates. The other problem is that the specimen will move out of focus, and therefore deviation in the Z coordinate must also be accounted for.

We show that eucentric height is a factor in both of these problems. We address Z movement using the pre-calibration method described in [1]. We track X and Y movement using proportional control — i.e. aligning each image to a previous image taken at an adjacent angle, and adjusting the microscope’s image shift deflectors to bring the specimen back to the center of view.

Eucentric height for a given specimen is defined as the Z coordinate of the stage where the center of rotation of the specimen coincides with the tilt axis of the stage. When the specimen is at this height, the stage is said to be at optimal eucentricity. This height minimizes the total lateral movement when the specimen is tilted over a range of angles. However, even when the specimen is at eucentric height, we show that movement in X,Y, and Z is still problematic.

Using proportional control to track X and Y movement at low and medium magnification is reliable due to a greater signal-to-noise ratio compared to high magnification, and therefore has a lower chance of observing faulty image correlations. The method is simple; at every tilt angle the specimen image is correlated with a previous image taken at an angle closest to the current angle. For small step sizes (*<*4 deg) and with an appropriate stretch of one image, the image features are nearly the same and will not affect correlation. The displacement between these two images is used to update the microscope’s image shift action using SerialEM’s AlignTo function. Overall, this method is fast and has negligible error. Most notably, time is saved by avoiding off-target tracking shots, which are more robust but slower.

Z movement is estimated by fitting calibrated data to a model. Focus measurements are taken at a nearby location along the tilt axis to estimate a sinusoid model of the Z coordinate as a function of tilt angle. In practice, we find that calibrating every time before collecting a tilt series is a good trade-off between runtime and accuracy compared to methods which calibrate less frequently. As mentioned in [1], errors in setting eucentric height will vary each time a eucentricity routine is performed, which can lead to poor estimates of Z movement. Calibrating at each specimen location avoids this problem.

## II. Stage Behavior at Different Heights

This section illustrates how the stage moves in the Y and Z coordinates for several initial heights around eucentric height, and shows the effects of perturbation error when this height is not optimal.

### A. Movement in Z

Figure 1 shows the behavior of the stage in the Z coordinate for several height offsets from a nominal eucentric height. The procedure to obtain the data is as follows:

**Fig. 1.**
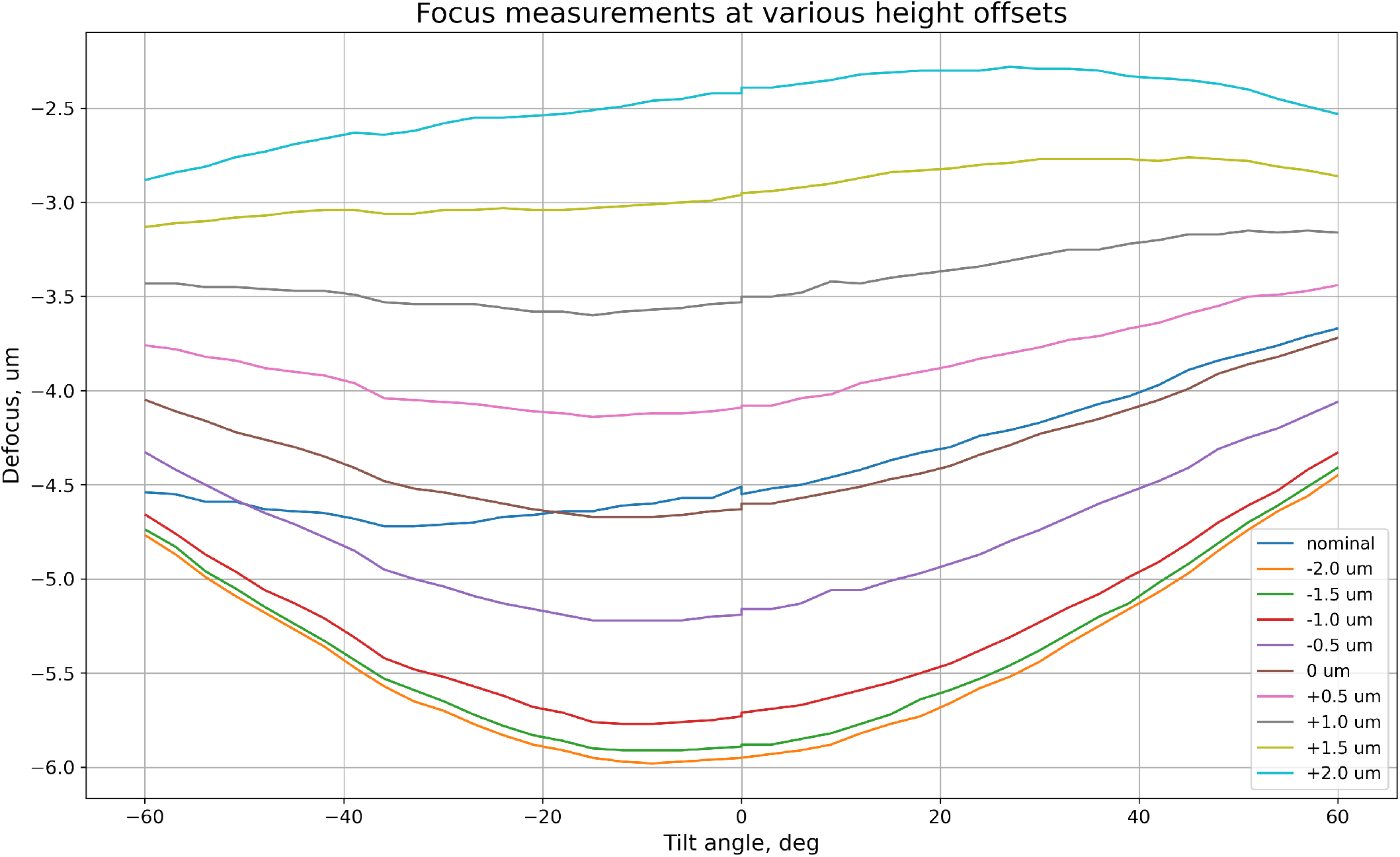
Focus measurements at different height offsets from a nominal eucentric height. Each tilt series was collected in a bidirectional scheme (angular range: 0 to*±*60 deg; step size: 3 deg). The series for the nominal eucentric height is inconsistent with the others, which shows that the behavior varies across routines that adjust stage height. The first two series at -2 um & -1.5 um offsets are inconsistently spaced due to stage backlash.

1. SerialEM’s fine eucentricity routine was performed to move the stage to a nominal eucentric height
2. A set of focus measurements were acquired in a bidirectional tilt series scheme, using image shift to keep the specimen point centered
3. The stage was lowered by 2 um
4. Focus measurements were similarly acquired in increments of 0.5 um for 9 iterations

The pre-calibration method is used to predict the Z coordinate. The method takes a sample of 5 focus measurements (at -40, -22, 0, 22, 40 deg) at a nearby area along the tilt axis, and does a least squares fit to the following sinusoid model

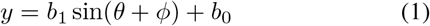

with parameters (*b*_1_, *b*_0_, *ϕ*), where *y* is the measured focus at the angle *θ*. This model encapsulates the rotation of an object around an ideal tilt axis, with inital displacements both laterally and in height.

A limitation of the pre-calibration method described above is that the specimen point cannot be centered due to large feature differences at different views. This means that the sample measurements are taken at different positions. Assuming that the nearby area around the specimen is planar, this modeling error is equal to *h* sin(*θ*) tan(*θ*), where *h* is the eucentric height error. For example, the error between the true focus and a sample measurement at 40 degrees and is 0.54*h*.

### B. Movement in Y

Figure 2 shows displacement in Y for several height offsets from a nominal eucentric height. The Y coordinate in SerialEM’s specimen coordinate system is perpendicular to the direction of the tilt axis. The displacements in Y from 0 degrees were acquired along with the focus measurements from Figure 1, by recording the image shift values that kept the specimen centered in the field of view. Even at near optimal heights, the maximum deviation is significant and requires tracking to ensure that the specimen stays in view. Movement in X (parallel to tilt axis) was also recorded but was insignificant and so is not shown.

**Fig. 2.**
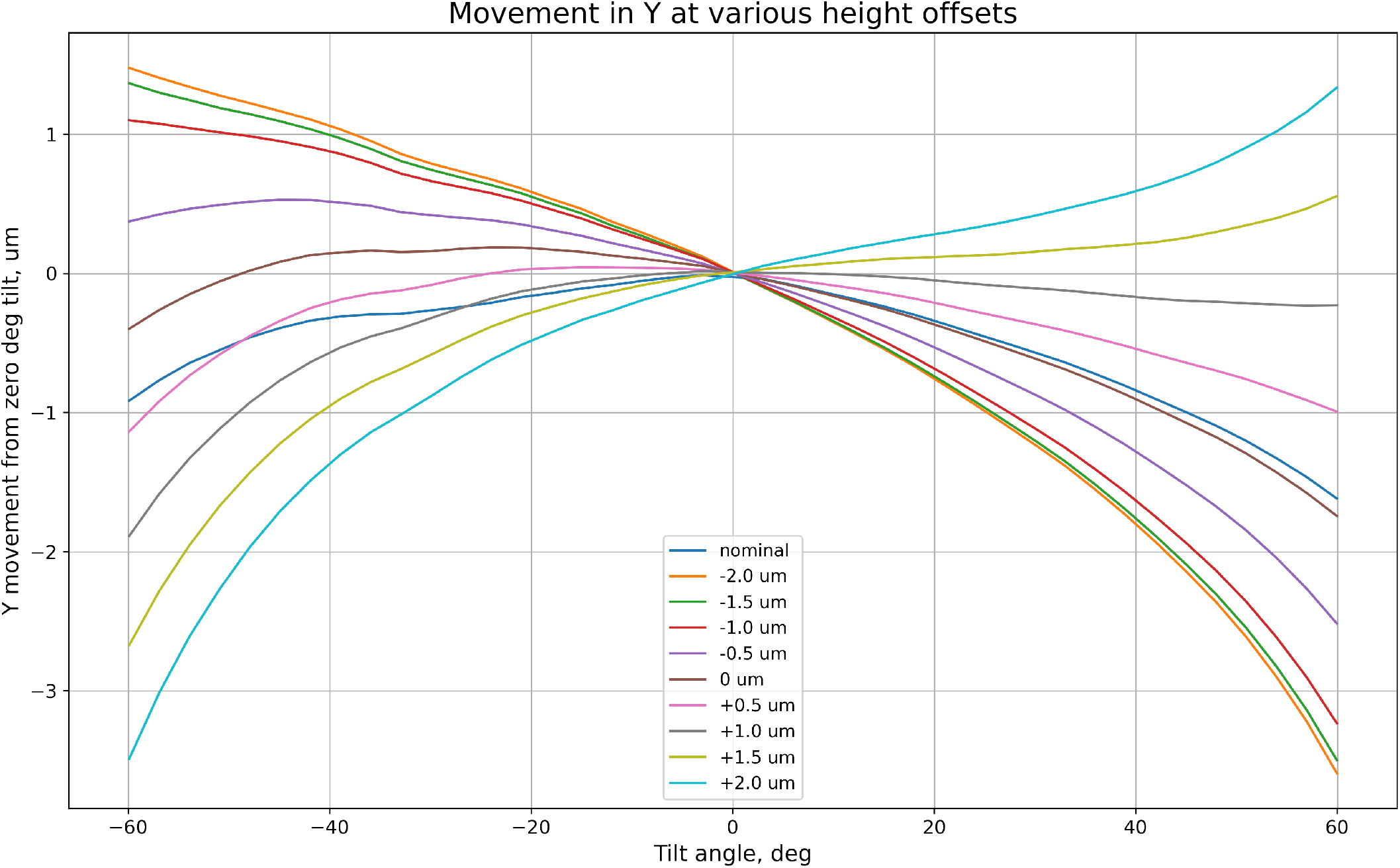
Y movement at different height offsets from a nominal eucentric height. The minimum deviation from 0 degrees out of all data series is roughly 1 um, which shows that tracking must be done even at eucentric height.

## III. Results

This section shows the effects of applying predictions for tracking Z movement, and a table of total runtimes used for several choices of tilt series parameters.

### A. Using predictions for Z movement

Figure 3 shows the effect of applying the model for predicting Z movement in an extreme example with particularly suboptimal eucentricity. Compensating for Z movement limits the maximum deviation to about 1 um from 0 deg tilt.

**Fig. 3.**
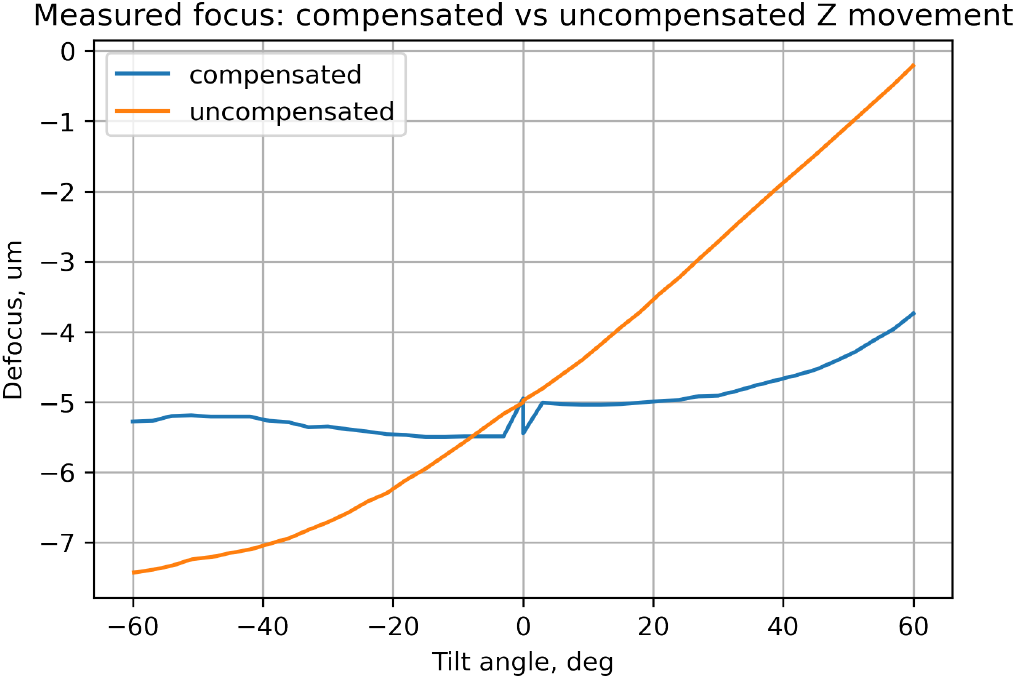
An extreme case of poor eucentricity. The uncompensated tilt series only tracks the specimen in X & Y without regarding Z. The compensated tilt series uses FastTomo. The measurements for compensated Z movement used the same bidirectional scheme as before (0 to -60 deg, 0 to +60 deg, 3 deg increments). Note: the uncompensated tilt series used a unidirectional scheme (−60 to +60 deg, 3 deg increments), so there is no discontinuity at 0 degrees.

### B. Runtime

Table I lists total runtimes for several choices of parameters. Each tilt series included the Z calibration routine, which on average takes 70 seconds.

**TABLE I.** Runtime

| Tilt series scheme | Angular range | Step size | Runtime |
| --- | --- | --- | --- |
| Unidirectional | -45 to +45 deg | 3 deg | 5 min 44s |
| Unidirectional | -45 to +45 deg | 2 deg | 7 min 8s |
| Bidirectional | 0 to $\pm 60$ deg | 3 deg | 7 min 12s |
| Bidirectional | 0 to $\pm 60$ deg | 2 deg | 9 min 16s |
| Dose-symmetric | 0 to $\pm 60$ deg | 3 deg, groups of 3 | 8 min 47s |
| Dose-symmetric | 0 to $\pm 60$ deg | 2 deg, groups of 4 | 11 min 4s |

Tested on an FEI Talos Arctica. Exposure time per image was 1.16s using a Gatan K3 camera.

## IV. Discussion

The choice of proportional control as a tracking method has several benefits, namely speed and simplicity. Without the expensive operation of separate tracking shots which alter beam and magnification settings, the majority of time spent in the script is occupied by stage settling and camera recording time. This method is very reliable for a sufficiently large field of view of at least 1 um. For higher magnifications, two main problems occur during data collection. Firstly, the displacement between images at successive angles becomes too large relative to the field of view, resulting in insufficient overlapping area and thus failed image correlation. This is especially noticeable when eucentricity is not optimal. Secondly, the lower signal-to-noise ratio results in statistically unreliable image correlations. In the low-dose regime of cryotomography, there is a higher probability of a faulty correlation, at which point the specimen will be lost from view.

Other methods for tracking lateral movement that were considered include: calibrating X & Y in the same way for Z; iterative polynomial fitting; and PID control – a widely popular feedback control algorithm used in industrial applications. In the calibration method, the lateral movement is predicted from a model and applied later. This method can be potentially unreliable because of the same concern for Z calibration, which is that setting eucentric height has varying error, even at the same stage location. An error in Z even as small as 0.5 um can have a lateral error of almost 1 um at maximum tilt, i.e. the sample will move out of view. Doing calibrations at every specimen location still did not alleviate this problem and added extra time to the calibration routine. A second method that was tested was iterative polynomial fitting. Unlike the calibration method, this method updates a polynomial fit to image displacements as data is collected, which is then used to make a prediction for the next displacement. This method does better, but still has about 0.5 um error at maximum tilt. Moreover, other considerations need to be made, such as robust regression to outlier data, which is observed on some microscope stages as large jumps at small nonzero angles. Another method that was tested was PID control, which is a historically well-known feedback controller which requires tuning gain parameters for the P, I, and D terms. Using very precise tuning, this method does slightly better than calibration and polynomial fitting, but was found to be impractical for the user to tune and thus unstable. The proportional control algorithm described in this paper does not require parameter tuning; in fact, it is a simple case of PID control with unity P gain and zero I & D gain. Ultimately, the simplicity and sufficient accuracy of proportional-only control is the reason why it was implemented.

The model for Z movement was chosen to be a sinusoid instead of a linear equation since the observed measurements in Figure 1 were nonlinear for all heights, including those near eucentric height, and especially for those farther away. Although [2] & [3] show linear behavior near eucentric height, in practice there is a precision error in setting eucentric height which is purportedly 0.1-0.25 um [1], causing the stage to move nonlinearly. Furthermore, the movement is not necessarily identical at different stage locations, even within the same grid square. Other models such as splines may also be effective, although we find that a sinusoid is accurate enough. In the case that the behavior is largely linear, it can be approximated reasonably well by *sin* over half of the tilt range, without the need to add a linear term to the model and risk overfitting.

## V. Conclusion

In this paper we showed how errors in setting eucentric height affect stage movement, and utilized proportional control and pre-calibration to simplify and speed up electron tomography data collection.

## VI. Code

The latest version of FastTomo can be found at https://github.com/alberttxu/FastTomo

## Acknowledgment

We would like to thank David Matronarde for insightful discussion, Jonah Murphy for general comments, and Jun Liu for feedback on user experience.

